# A programmed response precedes cell lysis and death in a mat-forming cyanobacterium

**DOI:** 10.1101/2022.10.17.512555

**Authors:** Jackie Zorz, Alexandre J. Paquette, Timber Gillis, Angela Kouris, Varada Khot, Cigdem Demirkaya, Hector De La Hoz Siegler, Marc Strous, Agasteswar Vadlamani

## Abstract

Cyanobacteria form dense multicellular communities that experience transient conditions in terms of access to light and oxygen. These systems are productive but also undergo substantial biomass turnover, supplementing heightened heterotrophic respiration and oxygen drawdown. Here we use metagenomics and metaproteomics to survey the cellular response of a mat-forming cyanobacterium undergoing mass cell lysis after exposure to dark and anoxic conditions. A lack of evidence for visral, bacterial, or eukaryotic antagonism contradicts commonly held beliefs on the causative agent for cyanobacterial death during dense growth. Instead, proteogenomics data indicated that lysis resulted from a genetically programmed response triggered by a failure to maintain osmotic pressure in the wake of severe energy limitation. Cyanobacterial DNA was rapidly degraded, yet cyanobacterial proteins remained abundant. A subset of proteins, including enzymes involved in amino acid metabolism, peptidases, toxin-antitoxin systems, and a potentially self-targeting CRISPR-Cas system, were upregulated upon lysis, indicating involvement in the programmed cell death response. We propose this natural form of programmed cell death could provide new pathways for controlling harmful algal blooms and for sustainable bioproduct production.

## Introduction

Eutrophication, resulting in expansion and prolongation of cyanobacterial mats and blooms, is expected to increase in frequency and severity with climate change and continuing industrialization [1, 2]. When these mats and blooms disintegrate, heterotrophic respiration of the released organic carbon can lead to extreme hypoxia, resulting in suffocation of fish and other marine sea life. Furthermore, many bloom-forming cyanobacteria release potent toxins upon lysis that have adverse health effects for humans and livestock [3]. Similarly, cyanobacteria grown for biotechnological purposes experience periodic mass culture crashes that negatively impact commercial operations [4]. Despite the economic and environmental importance, surprisingly little is known about the cause leading to the breakdown of dense cyanobacterial growth and the molecular response of cyanobacterial cells prior to their death [5, 6].

The main causes for cyanobacterial death include viral attack, predation, and programmed cell death [7]. Cyanobacterial viruses (cyanophages) and eukaryotic grazers are known to control cyanobacterial dynamics in natural freshwater and marine environments and have also been implicated in mass die-offs of cultures and blooms [8, 9]. Recently, however, more attention has been paid to the potential importance of programmed cell death in defining cyanobacterial dynamics in nature as a response to both biotic and abiotic threats [7, 10]. Programmed cell death in single-celled prokaryotes would appear to be an evolutionary paradox. However, evidence for this process has been observed in several situations, including thwarting viral propagation, and in multicellular communities, providing nutrients and metabolites to “persister” cells, which are members of a genetically analogous population that do not undergo cell death but rather persevere until conditions improve [11, 12]. Programmed cell death has been considerably less studied in prokaryotes compared to eukaryotes, but some hallmarks, including DNA fragmentation, protein degradation, degradation of ribosomes and RNA, and the involvement of reactive oxygen species are known [11]. Global molecular surveys of genes involved in prokaryotic programmed cell death are lacking, so the cell state leading to this process is largely unknown [6].

To investigate the process of cell death in cyanobacteria, we used an alkaliphilic cyanobacterial consortium that was observed to undergo a mass lytic event during a 12-day dark and anoxic incubation. This cyanobacterial consortium was previously isolated from the productive benthic mats of an alkaline (pH > 10, alkalinity > 0.5 M) soda lake [13, 14], and consists of the cyanobacterial species “*Candidatus* Phormidium alkaliphilum” [15] (renamed to *Sodalinema alkaliphilum* in GTDB version 214) in association with several other microbial members [16]. The dark and anoxic incubation could be considered similar to the conditions that the cyanobacteria would experience in nature by sinking into the aphotic regions of microbial mats or lake sediments [17, 18]. Interestingly, in the natural soda lake environment, the prolific growth of cyanobacteria in these mats does not appear to translate into a build-up of biomass or sediment. Instead, the presence of steep sulfide and oxygen gradients within the mats and lake sediments, indicates that cyanobacteria are rapidly turned over [17, 18].

We monitored the lysis of cyanobacterial cells by performing metagenomics and metaproteomics on samples collected throughout a 12-day dark and anoxic incubation of the consortium. From this data, there was no evidence for ecological interactions such as predation by other bacteria or viruses as the cause of cell lysis. Instead, proteogenomic data suggested that lysis resulted from programmed cell death provoked by energy starvation. We propose that mass die-offs and culture crashes commonly associated with dense growth of certain cyanobacterial taxa could in some cases be due to a similar genetically programmed response to adverse conditions.

## Results

### Cyanobacterial cell death accompanied by degradation of cyanobacterial DNA but persistence of cyanobacterial proteins

We subjected an alkaliphilic, mat-forming, cyanobacterial consortium to a 12-day dark and anoxic incubation, comparable to what it might experience upon sinking or sediment burial in its natural soda lake habitat. We collected samples from the solids (mat biomass) and supernatant (lysed and free-living) fractions every two days that were used for metagenomics and metaproteomics analysis. This data was then examined for signs of death-causing agents including viral attack, predation, and programmed cell death.

Initially, 72 % of the DNA extracted from the solid fraction originated from *Ca.* P alkaliphilum, but only 3.6 % of this DNA remained after 6-8 days in the dark. After 12 days, DNA from *Ca.* P alkaliphilum was barely detected (0.15%) (Fig. 1B). A similar pattern was observed in the supernatant fraction, as initially 20% of DNA could be attributed to *Ca.* P. alkaliphilum, which decreased to 1.6% by day 12 (Fig. 1C). In contrast, cyanobacterial proteins persisted throughout the incubation, always making up at least 65% of the protein composition in the solids fraction (Fig. 1E) and increasing to >80% of the supernatant fraction (Fig. 1F), suggesting a discrepancy in the way that the two biomolecules were degraded. Coinciding with the decrease of cyanobacterial DNA there was an increase in concentrations of fermentation products such as acetate and propionate (Fig. S1). Because acetate mainly accumulated before cyanobacterial lysis, *Ca.* P. alkaliphilum itself was likely responsible for its production [15, 19].

**Fig. 1.**
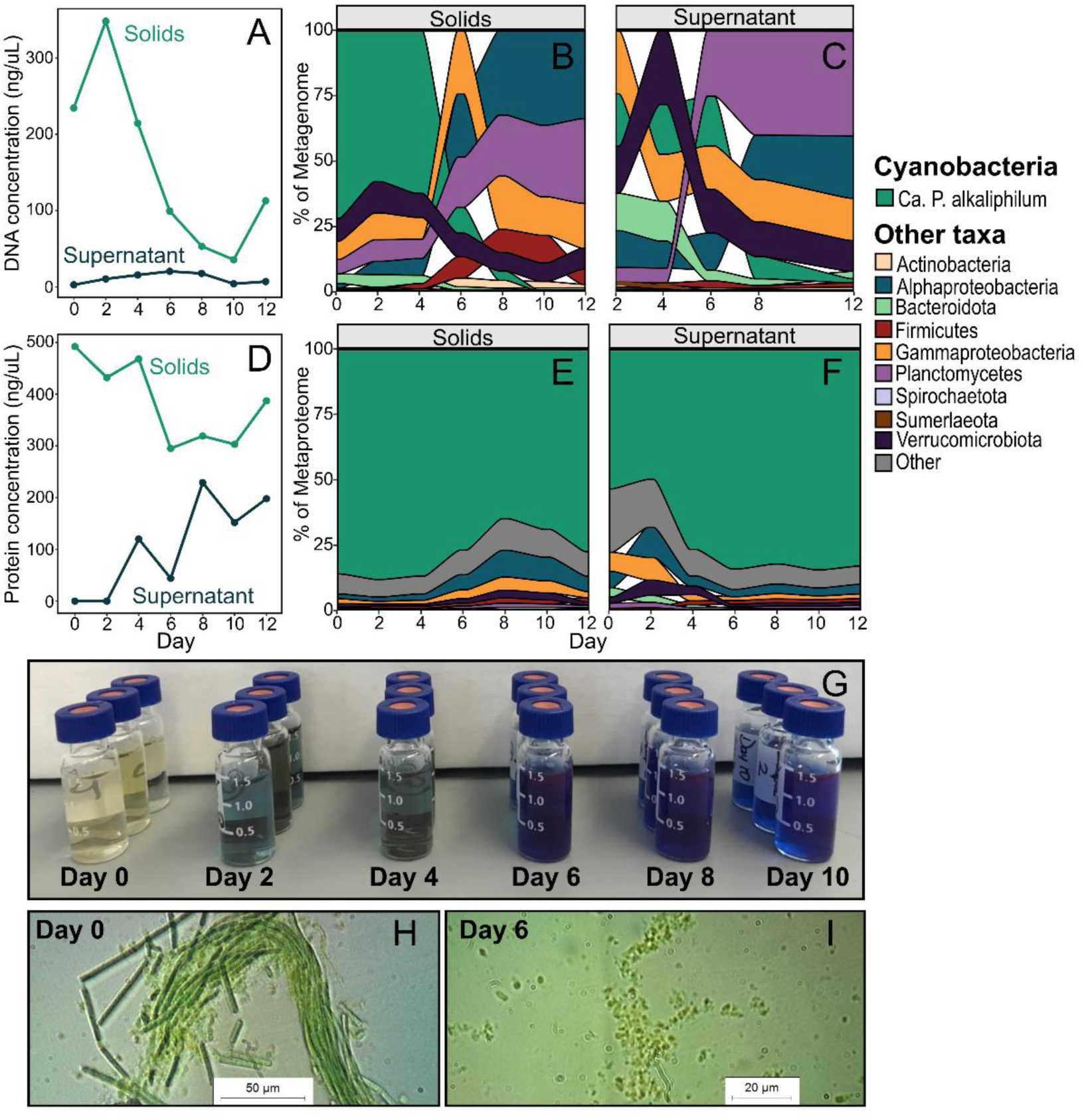
Observations during the 12-day dark and anoxic incubation. Concentration of DNA (**A**) and protein (D) in the solid and supernatant fractions. Microbial composition of consortium determined from the DNA in the solids (B), and supernatant (C) fractions, and from the protein content in the solids (E), and supernatant (F) fractions. Image showing the colour of the supernatant fraction of each sample taken during the incubation (G). Brightfield microscopy images of the cyanobacterial consortium taken on day 0 (H), and day 6 (I) of the incubation.

Propionate increased later in the incubation and was likely produced by other bacteria fermenting compounds within the cyanobacterial lysate. By day 6, the supernatant was coloured intensely blue (Fig. 1G) and contained a large amount of the internal antenna protein, phycocyanin, based on UV/Vis spectrometry (Fig. S2). Proteomics showed that phycocyanin made up 22%-32% of the protein in supernatant samples (Fig. S3). Microscopy showed that the cyanobacterial cells were lysing and breaking apart by the sixth day of incubation, explaining the presence of phycocyanin in the external medium (Fig. 1HI). Cell lysis of densely populated cyanobacterial blooms and the subsequent blue colour change caused by the release of phycocyanin is a phenomenon that has been observed previously in freshwater lakes, but the mechanism of these lysis events remains unknown [20, 21].

### Minimal evidence for cell death due to predation or viral attack

The possibility of predation by a larger eukaryotic cell or an antagonistic bacterial species was evaluated using the metagenomics data. Eukaryotic ciliate grazers affiliated with *Schmidingerothrix* were present in the consortium, however their abundance, calculated through copies of the rDNA gene in the metagenomes, was low (<2%) and did not increase over the course of the incubation (Fig. 2A), suggesting that eukaryotic predation was not directly responsible for the collapse of the cyanobacterial population.

**Fig 2.**
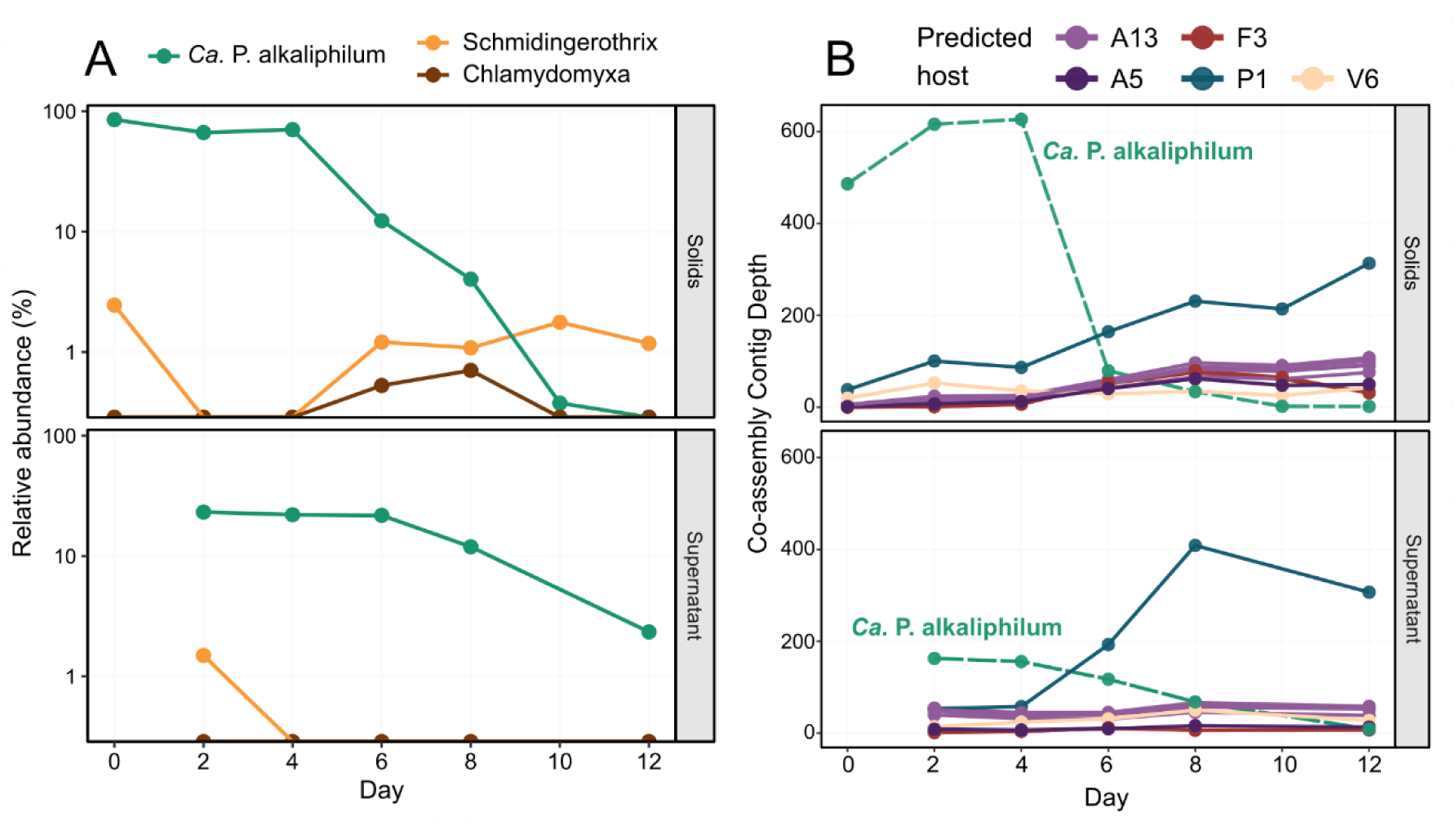
Dynamics of protist and viral sequences over the incubation. Relative abundance of *Ca.* P. alkaliphilum (green) and the two eukaryotic species identified (orange and brown), over the course of the incubation (A). Relative abundance based on rRNA genes retrieved from metagenomes using PhyloFlash. Y-axis is presented as a log-scale. Abundance of viral associated contigs over the incubation (B). The figure displays the top ten most abundant contigs with potential viral association. The top ten viral contigs had high confidence BLAST hits (>98% sequence identity) to contigs from MAGs, so the viral contigs are coloured based on their potential bacterial host. The taxonomy of the hosts are A13: Rhodobacteraceae, A5: *Pararhodobacter*, F3: *Alkalibacterium*, P1: UBA6054 (genus of Planctomycetota), V6: Verruco-01 (family of Verrucomicrobiota). The green dashed line shows the average contig depth for the *Ca.* P. alkaliphilum co-assembly MAG over the same samples.

Provisional genomes, or metagenome assembled genomes (MAGs), were acquired for the main cyanobacterial species, *Ca.* P. alkaliphilum, in addition to 59 other species from eight bacterial phyla including Proteobacteria, Bacteroidota, Firmicutes, Planctomycetota and Verrucomicrobiota (Fig. 1, Table S1). The observed bacterial dynamics, and the diverse and enhanced expression of carbohydrate active transporters, like the TonB-dependent transporters (expressed by at least 24 different MAGs), are akin to the succession of bacterial populations after phytoplankton blooms in other aquatic systems [22]. However, although the relative abundance of heterotrophs increased during the incubation, the abundances of heterotrophs relative to each other changed only modestly (Fig. 1) and no single species benefited in proportion to the magnitude of the cyanobacterial decline. Total DNA concentrations decreased (Fig. 1A), released cyanobacterial proteins persisted and fermentation (producing acetate) declined after the lysis of the cyanobacteria (Fig. S1). This might mean that the mainly aerobic consortium members did not have time to consume the released proteinaceous cyanobacterial lysate anaerobically. Taken together, our data did not point at a role of antagonistic bacteria in the cell lysis event.

Cyanophages, viruses that specifically target cyanobacteria, play an important role in the fate of cyanobacteria in natural environments, and consequently in global carbon and nutrient cycles [23, 24]. *Ca.* P. alkaliphilum contains seven CRISPR arrays, suggesting previous viral infections [25]. However, no viral proteins were identified in the metaproteome, and no viral contigs with regions of sequence identity to the *Ca.* P. alkaliphilum genome were identified in the assembly [26] (Fig. 2B). The most abundant contig of viral origin identified by *Virsorter* [27] was inferred to be a prophage of a Planctomycetota, and not associated with *Ca.* P. alkaliphilum. It also only reached 50% of the average cyanobacterial pre-collapse sequencing depth (Fig. 2B). In the event of a viral-mediated lysis, the depth of the associated viral contigs would be expected to increase at least 10-fold (burst size) in comparison to the depth of the cyanobacterial host contigs [28]. The comparatively low abundance of viral associated contigs indicated that a mass viral-induced lysis of the cyanobacterial cells did not occur.

### Extensive changes in cyanobacterial proteome allocations pre and post lysis suggest genetic programming led to cell death

Finally, we explored if a genetically programmed signal was the most likely cause of cyanobacterial lysis. Total cyanobacterial protein abundance remained relatively unchanged during the 12-day incubation, and included detection of just over 2,000 proteins, accounting for 52% of the predicted proteome of *Ca.* P. alkaliphilum (Table S2). Of the expressed proteins, 459 increased by at least two-fold between the beginning and the end of the dark incubation, while 1,039 proteins decreased expression over 50%. In general, cyanobacteria do not drastically change their proteome composition in response to diel cycling [29, 30]. Thus, a greater than twofold change in approximately 75% of the expressed proteins suggests that *Ca.* P. alkaliphilum had mounted a stress response that was outside the normal range of proteomic circadian cycles.

To better understand protein dynamics, we conducted a point biserial correlation analysis. This analysis identified proteins that were statistically linked to different time points and sample fractions during the incubations. The groups investigated in detail were the pre-lysis solids fraction (living stressed cells), the post-lysis solids fraction (surviving cells and some lysed cells), and the post-lysis supernatant fraction (lysed cells) (Fig. 3, Table S3). We also identified proteins associated with both the pre-lysis solids and the post-lysis supernatant (Fig. S4). This group likely contained intracellular proteins released into the media during lysis. In the following sections, we will provide more details about these findings.

**Figure 3.**
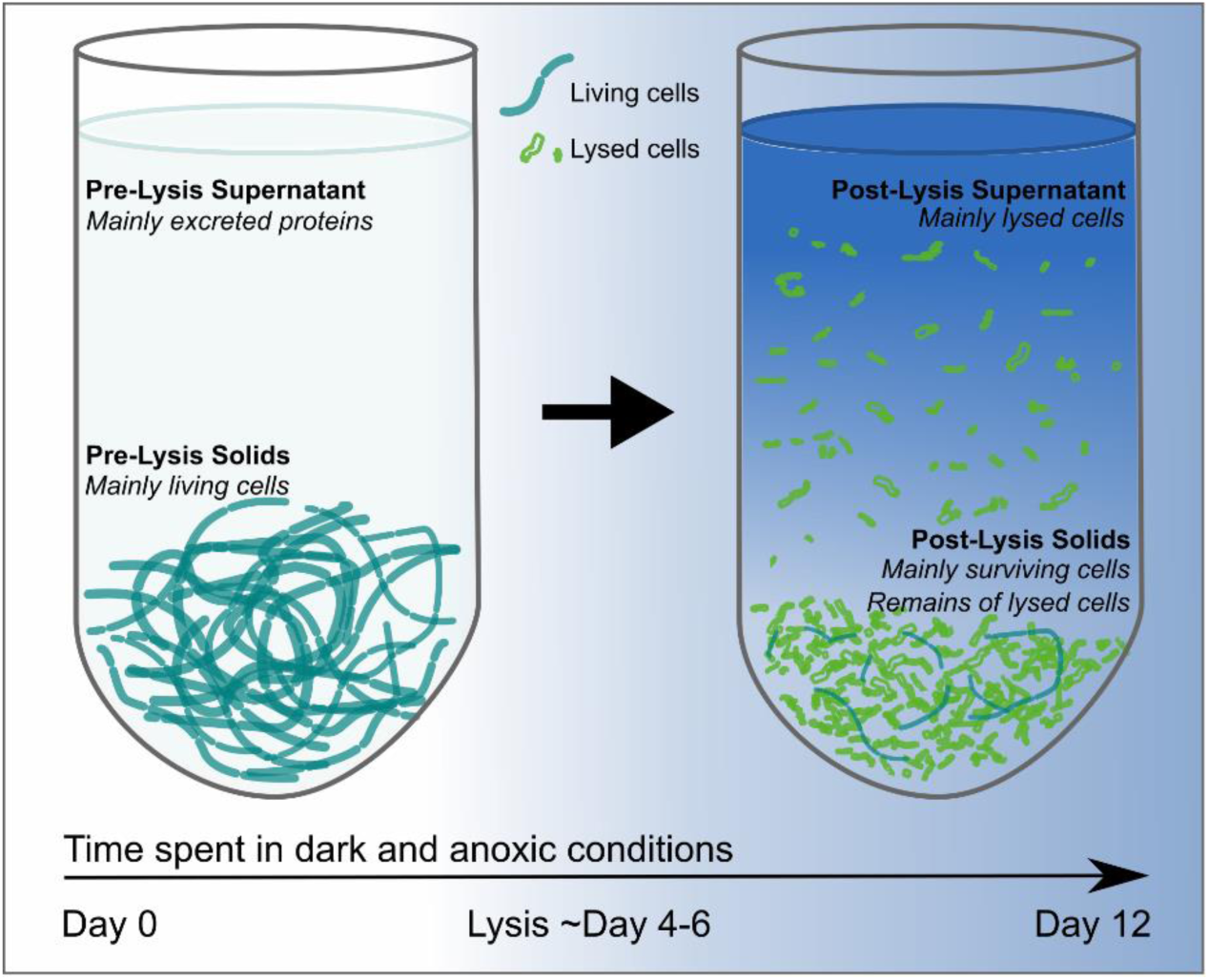
Schematic showing the expected cyanobacterial content in each fraction tested in the point biserial correlation analysis.

There were 52 proteins significantly more abundant in the solids fraction post-lysis. These proteins could have been upregulated in response to stress, or they could represent the proteome of surviving persister cells. The proteins upregulated in the solids fraction post-lysis included subunits of ATP synthase, cytochrome b6f, Photosystem I (PSI), and Photosynthetic Complex I (formerly called NDH-I). The latter is a homolog of Respiratory Complex I and is known to form a supercomplex with PSI [31]. These changes are signs of a shift from linear electron flow to cyclic electron flow around PSI [32, 33] (Table S2). In cyclic electron flow, electrons are cycled around PSI via passage through cytochrome b6f complex, ferredoxin and Photosynthetic Complex I [34, 35, 36, 37]. This results in the formation of a proton gradient to generate ATP, but no formation of NADPH. In cyanobacteria, cyclic electron flow is favoured when the cell is in an overly reduced state, and the ratio of ATP/NADPH is low [38]. During the incubation, the relative expression of PS1 increased nearly four-fold to 12.7% of the metaproteome by day 12. On this day, it was three-fold higher than the expression of Photosystem II (PSII), in contrast to the start of the experiment where PSI and II were expressed at an equal ratio. Proteins not required for cyclic electron flow, like ferredoxin-NADP+ reductase and the oxygen evolving complex of PSII decreased ten and two-fold, respectively, during the same period.

Previous studies have identified an increase in cyclic electron flow in response to dark and anoxic conditions and have hypothesized that it could be a defensive strategy in preparation for the return of daylight and resumption of photosynthesis [39], or alternatively as a mechanism to jumpstart metabolism through the rapid generation of ATP once light energy returns [40]. In our incubation, light did not return, and the mounted stress response was ultimately unsuccessful.

Proteins that were statistically more abundant in the pre-lysis solids fraction, and downregulated before cell lysis, were largely involved in translation and transcription and included ribosomal proteins, elongation factors, translation initiation factors, tRNA ligases, and RNA and DNA polymerases (Fig. 4A). Additionally, proteins expressed more in the pre-lysis solids fraction included protein chaperones (e.g., GroEL), and transcriptional regulators, most with poorly defined functions and targets (Fig. 4A). During the incubation, the abundance of these proteins decreased significantly, by as much as 45-fold, and there was no corresponding increase observed in the post-lysis supernatant fraction. This indicates that these proteins were actively degraded rather than being released into the surrounding media during lysis (Fig. S4).

**Fig 4.**
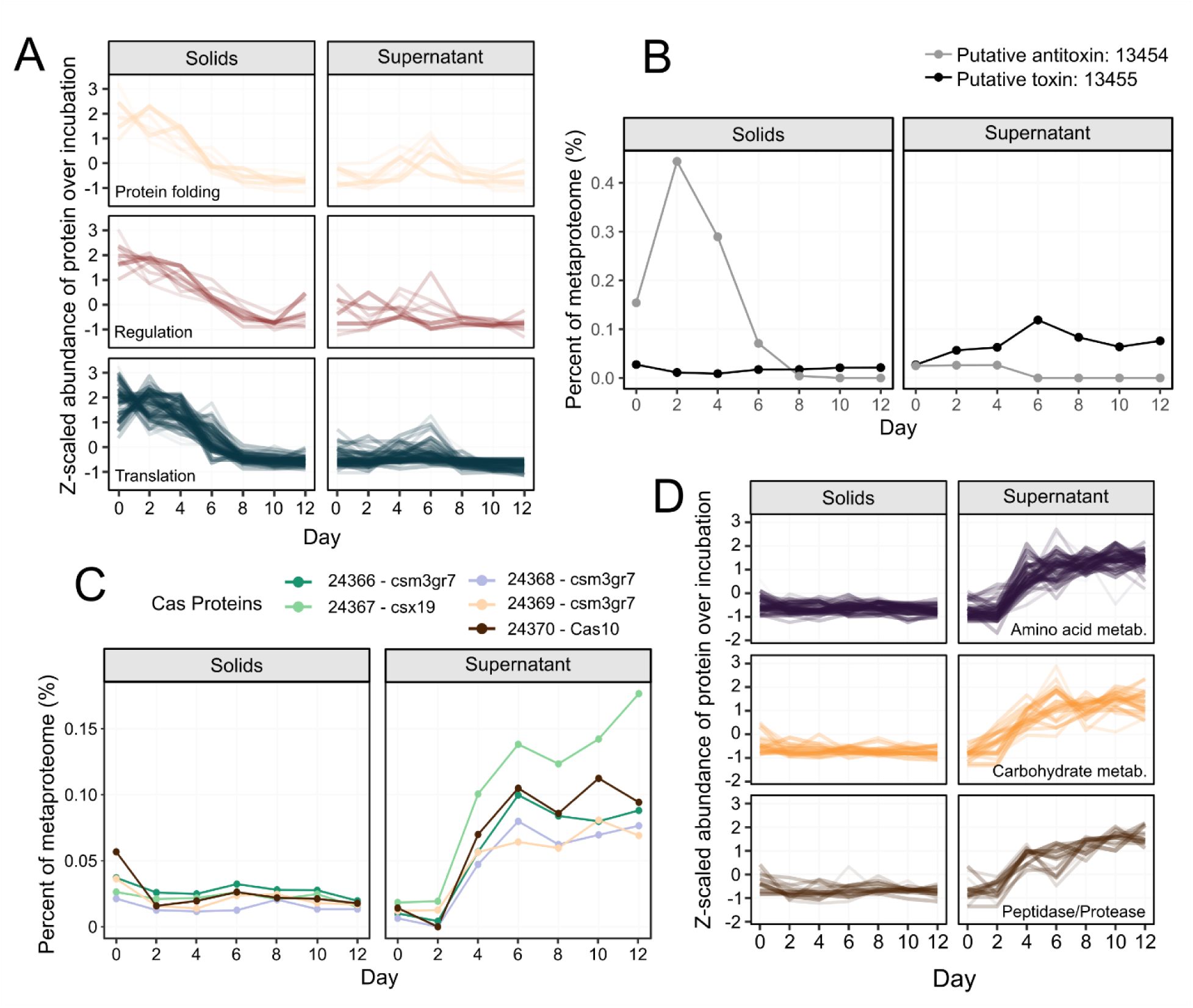
Dynamics of *Ca.* P. alkaliphilum proteins over the course of the incubation. Proteins from functional categories that decreased in abundance in the solids fraction leading up to cell lysis (A). Protein abundances were scaled across samples for visualization purposes. For a complete list of these proteins see Table S3. Metaproteome abundance of the potential toxin-antitoxin protein pair over the incubation (B). The numbers indicate the protein accessions (Data S1). Abundance of proteins located in a CRISPR-Cas operon increased in the supernatant post-lysis (C). Numbers correspond to protein accession (Data S1). Proteins from functional categories that were significantly higher in abundance in the post-lysis supernatant fraction (D). Protein abundances were scaled across samples for visualization purposes. For a complete list of these proteins see Table S3.

Within the *Ca*. P. alkaliphilum genome, there are multiple toxin-antitoxin systems [15]. One particular putative antitoxin protein (accession: 13454) was observed to have higher abundance in the pre-lysis solids fraction and was undetectable in the post-lysis incubation (Figure 4B). This protein shares sequence similarity with the transcriptional regulator AbrB [41, 42] and the MazE antitoxin [43]. Both of these proteins are known to bind and inhibit the transcription of toxin genes [44, 45]. The specific toxin associated with this putative system remains unknown, but it is possible that the protein directly downstream (accession: 13455) could fulfil the role, as it possesses a ribonuclease domain and showed significantly higher abundance in post-lysis supernatant samples (Figure 4B). In MazEF toxin-antitoxin systems, the toxin MazF often functions as an endoribonuclease [43], exerting its toxicity by inhibiting protein synthesis [46, 47, 48]. MazF has also been implicated in the selection of cells that undergo cell death versus those that survive under stressful conditions [49]. It is important to note that this association is speculative as the study of toxin-antitoxin systems in prokaryotes is still relatively new and has primarily focused on model organisms [50], leading to a potential underestimation of the full diversity of these systems.

In conducting this differential abundance analysis, it was important to distinguish between intracellular proteins that are naturally released into the media during the lysis process and proteins that were statistically more abundant in the post-lysis supernatant fraction, indicating upregulation in lysing cells. We differentiated these two groups by distinguishing proteins that were enriched in both the solid pre-lysis and supernatant post-lysis fractions (intracellular proteins) from proteins that were enriched exclusively in the supernatant post-lysis fraction (potentially involved in lysis). There were 16 proteins that showed significant association with both the pre-lysis solids and post-lysis supernatant fractions. These proteins primarily included antenna-complex proteins such as phycobilisome linker proteins and allophycocyanin, as well as proteins involved in carbon metabolism like RuBisCO (rbcL), phosphoglycerate kinase, and transketolase (Fig. S4).

A total of 142 proteins showed significant increase in abundance exclusively in the post-lysis supernatant fraction. These proteins were likely upregulated in cyanobacterial cells shortly before their death and subsequently released into the surrounding media during lysis. As a result, it is plausible that these proteins played a role in causing cyanobacterial death. A highly upregulated group of proteins included five CRISPR-associated (Cas) proteins forming a Type IIIA CRISPR-Cas operon (accessions 24366-24370). Expression of these Cas proteins increased over 10-fold between pre- and post-lysis in the supernatant (Fig 4C). Because we were unable to find evidence for viral lysis of *Ca*. P alkaliphilum (Fig. 2B), it seems possible that this Cas response was independent of viral or immune activity and may instead be associated with the cyanobacterial response to stress. In support of this theory, previous research has suggested that ancestral CRISPR-Cas effectors were stress response systems that triggered programmed cell death after activation by a signalling molecule [51, 52]. CRISPR Type IIIA effector complexes consist of a Cas10 protein and other subunit proteins Csm3 (Cas7), and Csx19 [53] acting as a multi-subunit nuclease [54]. The nuclease activity of the Type IIIA systems has previously been linked to cell death and dormancy [55]. Furthermore, CRISPR systems have also been shown to work in association with toxin-antitoxin systems [56, 57, 58, 59, 60].

Proteins associated with the supernatant post-lysis fraction also included several peptidases and proteases, as well as over 30 proteins involved in amino acid metabolism (Fig 4D). This indicates active proteome remodelling or degradation. Additionally, several proteins involved in carbohydrate degradation and synthesis, as well as electron and iron carriers such as ferredoxin and bacterioferritin were enriched in the supernatant post-lysis.

### Evidence for similar cellular lysis event found in source soda lake environment and in commercial Arthrospira platensis culture

Evidence for cyanobacterial lysis occurring *in situ* was investigated using 30 cm cores of cyanobacterial mats and underlying sediment from the culture source, alkaline Lake Goodenough (Canada) (Fig. 5A). 16S rRNA gene amplicon sequencing of sectioned cores showed high abundance of cyanobacteria at the top of the mat (Fig. 5BC). The abundance of cyanobacteria like *Phormidium* and *Nodosilinea* decreased rapidly and became essentially negligible two cm below the sediment surface. Rapid turnover of cyanobacterial biomass explained the previously observed steep sulfide gradients [17]. Sulfide likely builds up below the mats after the depletion of oxygen because sulfur reducing bacteria oxidize fatty acids, hydrogen, and other cyanobacterial degradation products and reduce sulfate and other sulfur-compounds to sulfide. Amplicon sequencing showed the ecological success at depth of thiosulfate and elemental sulfur reducing *Dethiobacter* [61, 62], and the sulfate, thiosulfate, and sulfite reducing *Desulfonatronovibrio* [63] (Fig. 5DE).

**Fig. 5.**
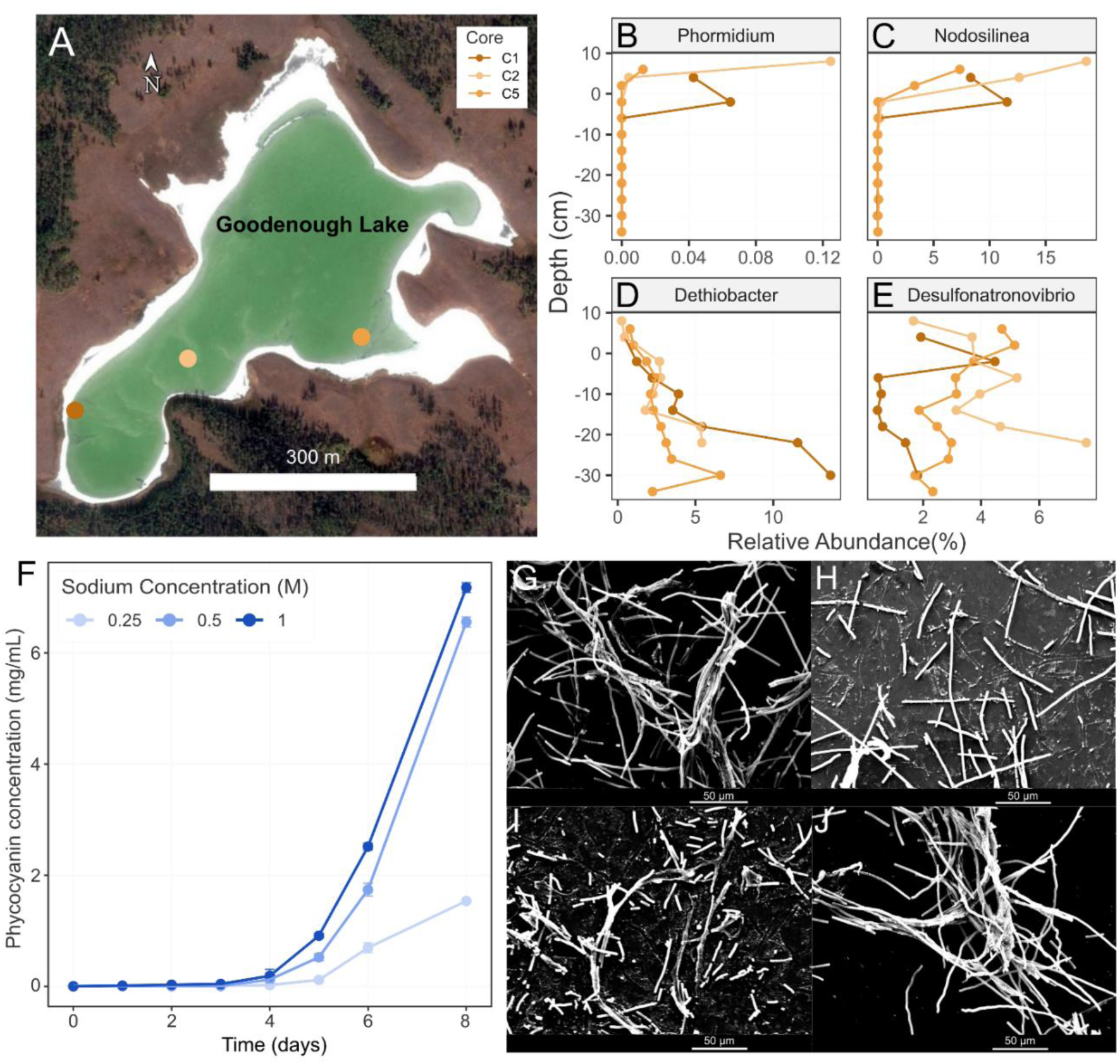
Data from sediment cores and experiments varying sodium concentration. (**A**) Map of sampling locations within Goodenough Lake, British Columbia, Canada. Distribution of genera from 16S rRNA gene abundance: (B) Cyanobacteria *Phormidium* (*Ca*. P. alkaliphilum), (C) Cyanobacteria *Nodosilinea*, (D) Sulfidogenic thiosulfate and elemental sulfate reducing bacteria *Dethiobacter*, (E) Sulfidogenic, sulfate, sulfite and thiosulfate reducing bacteria *Desulfonatronovibrio*. Positive depth values in B-E represent the cyanobacterial mat and negative centimetres represent distance below the sediment surface. (F) Phycocyanin concentration in the supernatant fraction of dark and anoxic incubations with varying sodium concentrations in the media. Electron micrographs of cyanobacterial cells on day 0 (G), and day 5 of dark and anoxic incubations, with the original 0.5 M Na^+^ media (H), 1 M Na^+^ media (I), and 0.25 M Na^+^ media (J).

If energy depletion was the main stress leading to initiation of the programmed cell death response and the corresponding cyanobacterial cell lysis, increasing salinity prior to a dark and anoxic incubation should result in an earlier lysis event. The cells would need to spend more maintenance energy to cope with the higher osmotic pressure and would consequently deplete their reserves and enter a stressed state sooner. This hypothesis was tested by performing separate dark and anoxic incubations of the cyanobacterial consortium at higher (1M Na^+^) and lower (0.25M Na^+^) salinity.

The dark and anoxic incubation at 0.25M Na^+^, resulted in a cyanobacterial lysis event that occurred later and was less complete, characterized by a much lower concentration of released phycocyanin (1.5 mg/mL), compared to the original incubation at 0.5M Na^+^ (6.6 mg/mL) (Fig. 5F). In the incubation with 1M Na^+^, cell lysis occurred sooner, by day 5 (Fig. 5I), and the final concentration of phycocyanin was high (7.2 mg/mL). These results support the hypothesis that cyanobacterial cell lysis in these dark and anoxic incubations is initiated by depleted energy reserves. Cells in an environment of higher salinity require more energy to maintain osmotic equilibrium, and thus deplete energy reserves faster.

Lastly, we demonstrated that this cellular lysis occurs in cyanobacterial species outside the *Phormidium* genus by performing a similar dark and anoxic incubation on the commercially important species *Arthrospira platensis*, recently renamed to *Limnospira platensis*. A culture of this cyanobacterium was incubated in dark and anoxic conditions with 1M Na^+^ and after 10 days of incubation, a similar phycocyanin release was observed signalling that an analogous lysis event had occurred (Fig. S5).

## Discussion

Despite the ecological and economic importance associated with biomass turnover in cyanobacterial systems, there is a general lack of understanding of what causes the collapse of dense cyanobacterial mats and blooms, and what occurs at the molecular level in these cells before death. Here we used metagenomics and metaproteomics to gain insights into the molecular state of the cyanobacterial cells prior to and post lysis. Based on this data, we concluded that the mass cell lysis event was caused by programmed cell death in response to severe energy limitation and osmotic stress.

Proteogenomics showed that the cells altered their proteome to combat energy stress brought about by darkness, by the rearrangement of protein complexes in the thylakoid membrane to favour cyclic electron flow. Fermentation of available endogenous carbohydrates, like glycogen, cyanophycin, or the osmolytes sucrose, glucosyl glycerol, and trehalose, would have initially provided energy for sustaining cellular integrity and resulted in the observed increase in acetate [64, 65] (Fig. S1). After four days, proteins involved in transcription and translation were severely diminished in the solids fraction, a signal of decreased metabolism and arrested growth. With continued darkness, the supply of endogenous carbohydrates and osmolytes was likely depleted [66]. The ensuing starvation and osmotic stress may have triggered a programmed cell death response, possibly through toxin-antitoxin systems and/or the expression of Type IIIA CRISPR associated proteins in an upregulated operon, resulting in lysis of the cyanobacterial cells.

A similar lysis phenomenon was previously observed in dark and anoxic incubations of a thermophilic cyanobacterium, *Oscillatoria terebriformis*, isolated from hot spring microbial mats [67]. In that experiment, cell survival could be prolonged by the addition of an exogenous carbohydrate source (fructose), and/or a reductant (e.g., sodium thioglycolate). This might be another example where dwindling energy stores cause the lysis of a cyanobacterium under dark and challenging conditions. The addition of fructose sustained cell survival by providing another substrate for cyanobacterial fermentation. Whereas the addition of reducing agents could have quenched reactive oxygen species (ROS) produced under stress. A link between ROS and programmed cell death in both eukaryotic and prokaryotic cells has long been known [11, 68, 69], and the production of ROS has previously been associated with the activation of toxin-antitoxin systems upon stress [70, 71].

Evidence of this lytic bioprocess was found in the industrially cultivated species, *A. platensis*, as well as in cyanobacterial species from hot spring microbial mats [67], and the sediments of the original haloalkaline environment of *Ca.* P. alkaliphilum, suggesting that this phenomenon could be widespread among cyanobacteria in both engineered and natural systems. Ultimately, this may even open new avenues to control harmful algal blooms. Detection of free phycocyanin in lakes after blooms [72] already indicates that the same process could be relevant and, with follow up research, manipulated. Furthermore, cyanobacterial cellular lysis due to dark and anoxic incubation provided a way to access the internal pigment phycocyanin without costly and energetically intensive mechanical disruption [73]. Adapting this process to an industrial setting could reduce the costs and energy associated with the production of phycocyanin and other internal compounds.

This study provides important insights into mechanisms of mass cell death in cyanobacterial mats and blooms. From this data, we show that cellular machinery related to protein synthesis is specifically degraded during a prolonged dark period and hypothesize that there are previously unidentified roles for toxin-antitoxin systems and Cas proteins in cyanobacterial cell death. Additionally, this process could be a promising target for new antibiotics or algicidal compounds and harnessed as a commercially sustainable way to produce bioproducts.

## Materials and Methods

### Experimental setup and sampling

An alkaliphilic cyanobacterial consortium enrichment culture containing a single, abundant, cyanobacteria species, *Candidatus* Phormidium alkaliphilum (renamed to “*Sodalinema alkaliphilum*” in GTDB version 214) [13, 15, 16], was used as inoculum for the dark incubation. This consortium was originally sourced from alkaline soda lakes in the Cariboo Plateau region of Canada [14]. The cyanobacterial consortium was grown in continuous light (200 µmol photons/(m^2^·s)) in 10 L stirred glass vessels. The growth medium was previously described [74] and contained 0.5 M sodium (bi)carbonate alkalinity, at an initial pH of 10.3. After six days of photoautotrophic growth the culture was centrifuged for 30 minutes at 4,500 rpm to concentrate the biomass (Allegra X-22R, Beckman Coulter, USA). The wet biomass was then divided into 20 mL serum bottles sealed with butyl-rubber septa. Two grams of wet biomass were added to each serum bottle. The bottles were purged with N_2_ gas to create an anoxic headspace, and then placed at room temperature (21°C) in the dark. At 0, 2, 4, 6, 8, 10, and 12 days after the start of the incubation, two sacrificial samples were taken. To each sample, 5 mL of pH 7, phosphate-buffered saline solution was added and then the sample was centrifuged for 10 min at 4500 rpm, to separate biomass and supernatant.

For biomass pellets, the ash free dry weight of each sample was measured using NREL laboratory analytical procedures protocol [75]. For the supernatant, the concentration of the organic acids succinate, formate, propionate, butyrate, and lactate were measured using an UltiMate 3000 HPLC system (ThermoFisher Scientific, USA) equipped with an Aminex HPX-87H column and a UV detector, as previously described [76]. The phycocyanin and total protein concentration in the supernatants were measured as absorption at 620 nm and 280 nm respectively [77] using an Evolution 260 Bio UV-Visible Spectrophotometer (ThermoFisher Scientific, USA), with a standard curve prepared from laboratory-grade phycocyanin (Sigma-Aldrich, USA). Bright-field microscope images were taken using a Zeiss Axio Imager A2 Microscope (Carl Zeiss AG, Germany).

### DNA extraction

DNA was extracted directly from soda lake sediment and incubation solid samples using the Fast DNA Extraction Kit for Soil (MP Biomedicals, USA). For incubation supernatant samples, 250 µL of sample was used for each extraction. The extraction protocol of the manufacturer was followed, but additional purification steps were performed with 5.5 M guanidine thiocyanate [13]. For supernatant samples, the elution buffer was heated to 50°C prior to the elution step to increase yield. Still, supernatant samples from Day 0 and Day 10 did not yield enough DNA for metagenome analysis.

### 16S rRNA gene PCR and sequencing

Amplicon sequencing and library preparation of the DNA sediment samples was performed as previously described [13] using primer sets 926wF (5′-AAACTYAAAKGAATTGRCGG3′) and 1392R (5′-ACGGGCGGTGTGTRC3′) targeting bacteria [78, 79]. Prepared libraries were sequenced on the MiSeq Personal Sequencer (Illumina, USA) using the 2 × 300 bp MiSeq Reagent Kit v3. Amplicon sequencing results were processed using MetaAmp Version 3.0 [80], and the Silva database version 132 [81]. Paired-end reads were merged if they had less than eight mismatches in the overlap region and an overlap of >100 bp [13, 82]. The merged reads were further filtered by removing reads that were missing the forward or reverse primer and had more than one mismatch in the primer region. All reads were trimmed to a final of 350 bp and clustered into operational taxonomic units (OTUs) of >97% sequence identity [13, 82].

### Library preparation and metagenome sequencing

All biomass samples, and supernatant samples from days 2, 4, 6, 8 and 12 were prepared for metagenomic sequencing as previously described [14]. Briefly, DNA was sheared to fragments of ∼300 bp and libraries were created using the NEBNext Ultra DNA Library Prep Kit (New England Biolabs, Ipswich, MA). An Illumina NextSeq 500 sequencer (Illumina, San Diego, CA) was used for sequencing using a 300 cycle (2×150bp) high-output sequencing kit at the Center for Health Genomics and Informatics in the Cumming School of Medicine, University of Calgary, Canada.

### Metagenome Assembly and Binning

Raw, paired-end Illumina reads were filtered for quality using BBDuk (https://jgi.doe.gov/data-and-tools/bbtools/). Quality control consisted of trimming reads to 150 bp, trimming off adapter sequences, filtering out contaminants, such as the PhiX adapter, and clipping off low quality ends, all as previously described [14]. Paired-end reads from each sample were then merged with BBMerge [83]. Separate assemblies of the reads from each sample were performed using metaSPAdes version 3.12.0 with default parameters [84]. To increase binning success, one large co-assembly using the unmerged reads from all samples was conducted using MegaHit v1.2.2 [85]. Only contigs greater than 500 bp in length were processed further. The MetaErg pipeline [86] was used for prediction and annotation of genetic elements on each assembled contig.

Binning of assembled reads into metagenome-assembled-genomes (MAGs) was completed using MetaBat2 version 2.12.1 [87]. The binning step was performed on each sample’s assembly separately as well as the co-assembly. To generate sequencing depth data for binning, quality-controlled reads of each sample were mapped to the assembly of each sample using BBMap v38.84 (https://sourceforge.net/projects/bbmap/). Mapping results were summarized using the script, “*jgi_summarize_bam_contig_depths*”, part of the MetaBat package [88]. After binning, the program dRep [89] in conjunction with CheckM v1.0.11 [90] was used to determine the best (highest estimated completeness, and lowest estimated contamination) MAGs associated with each population. In total, 60 MAGs (>80% completeness, and <5% contamination) were identified for further processing and analysis. The program, gtdbtk v0.3.2 was used for the taxonomic assignment of each MAG [91].

The relative abundance of individual MAGs in each metagenome was calculated by mapping quality controlled raw reads from each sample onto the contigs of each MAG as well as the dereplicated contigs that remained unbinned. Again, BBMap (minid = 0.98) was used for this. Unbinned contigs were dereplicated using cd-hit-2d [92]. In this step, all contigs sharing > 90% sequence identity with a binned contig were eliminated. The number of reads that mapped to each contig was counted, and then the total counts for each contig of each MAG were summarized. To determine relative abundance, counts were normalized to MAG genome size and the number of mapped reads per sample. For the unbinned contigs, reads were normalized to the number of base pairs in all dereplicated unbinned contigs.

The program phyloFlash v3.3 (*Emirge* assembly) was used to obtain full length 16S and 18S ribosomal RNA (rRNA) gene sequences and their sequencing depth from the metagenomes [93]. The sequencing depths of rRNA sequences were used primarily to determine the population dynamics of species that did not assemble or form MAGs well, mainly eukaryotic protists.

### Analysis of viral contigs

Contigs potentially associated with viruses were identified from the metagenome co-assembly using VirSorter v1.0.6 [27]. BLASTn was then used to match the DNA sequence of viral contigs to the 60 MAGs. CRISPR arrays in the cyanobacterial MAG were identified from the MetaErg output [86], with the program MinCED (github.com/ctSkennerton/minced).

### Protein Extraction and LC-MS/MS mass spectrometry

Protein was extracted from biomass and supernatant samples as previously described [94], using the filter aided sample preparation (FASP) protocol [95]. To lyse cells, samples were added to lysing matrix E bead tubes (MP Biomedicals, USA) with SDT-lysis buffer (0.1 M DTT) in a 1:10 sample to buffer ratio. The tubes were then subjected to bead-beating in an OMNI Bead Ruptor (Omni International, USA) 24 for 45 s at 6 m s^−1^. For supernatant samples, 500 µL of supernatant was used for lysis. Supernatant samples from days 0 and 2 had low yields and so lysate was concentrated prior to protein extraction.

Peptides were separated by an UltiMateTM 3000 RSLC nano Liquid Chromatograph (Thermo Fisher Scientific, USA), using a 75cm x 75µm analytical column and analyzed in a Qexactive Plus hybrid quadrupole-Orbitrap mass spectrometer (Thermo Fisher Scientific, USA) as previously described [96]. A total of 2,000 ng of peptide was loaded, and each sample was run for 4 hours.

### Metaproteomics data analysis

The database used for protein identification was manually created using the predicted and annotated proteins from the binned and unbinned metagenomic sequences. Cd-hit was used to remove redundant sequences from the database using an identity threshold of 95% [92], giving preference to sequences that came from MAGs. Sequences of common contaminating proteins were added to the final database (http://www.thegpm.org/crap/). The final database contained 454,164 proteins. For protein identification MS/MS spectra were searched against the database using the Sequest HT node in Proteome Discoverer version 2.2.0.388 (Thermo Fisher Scientific, USA) as described previously [97]. Only proteins with one unique peptide, and with a protein false discovery rate (FDR) confidence of at least a level of “medium”, were kept for further analysis.

Relative protein abundances were estimated using the normalized spectral abundance factor (NSAF) [98]. MAG abundance in the metaproteome was estimated by adding the NSAF abundance of all proteins belonging to that MAG. In total, 3,286,730 MS/MS spectra were obtained, yielding 632,137 peptide spectral matches (PSMs), which corresponded to 10,408 expressed proteins after quality control.

To further investigate the protein expression dynamics during the dark and anoxic incubation, the *Ca.* P. alkaliphilum proteomics data was divided into pre and post lysis groupings. The supernatant pre lysis grouping included days 0 and 2, and the supernatant post lysis grouping included days 4-12. The solids pre lysis grouping included days 0, 2, and 4, and the solids post lysis grouping included days 6-12. These groupings were chosen based on NMDS clustering (Fig. S6) of the cyanobacterial proteomics data and the phycocyanin concentration in the supernatant (Fig. S2), which was used as a proxy for cell lysis. NMDS was performed using the metaMDS function from the R package *vegan* [99] based on Bray-Curtis dissimilarity of proteins with greater than 50 PSMs. Proteins that were statistically differentially abundant between these groupings from the *Ca.* P. alkaliphilum MAG were identified using point biserial correlation implemented with the *multipatt* function from the *indicspecies* package in R [100] (Table S3). The *multipatt* function was run with the “r.g” function and 9999 permutations. Only proteins that had at least 50 PSMs in total were included in the analysis, and a p-value of 0.05 was needed to be considered significant. The functional annotations of statistically significant proteins were further validated using BLASTP searches against the NCBI nr database.

### Sediment sample collection and preparation

Duplicate sediment cores were collected in April 2019 from Goodenough Lake (51.330°N, 121.64°W). The sediment cores were taken from 3 different locations within each lake (Fig. 1) using a 1.5-m single-drive Griffith corer from LacCore: National Lacustrine Core Facility (University of Minnesota). The sediment cores ranged in length from 25-50 cm. To reduce the mixing of water and upper sediment layers in the cores, Zorbitrol was used as a gelling agent to stabilize the sediment-water interface during transport. Cores were then stored upright at -20 °C.

Cores were removed from the -20 °C freezer and defrosted at room temperature (22 °C) for 2 hours. Cores were then horizontally sliced into 2 cm disks using a Dremel Multi-Max MM50 oscillating saw (Dremel, USA) at the lowest speed, used to reduce blade contact with the sediment. The blade was sterilized with 70% ethanol before each core section was sampled. To avoid the potential risk of contamination from the core liner or during sectioning, sediment in contact with the core liner was removed and the inner core was transferred to a 50 mL tube, sealed, and stored at -20 °C. The sediment from each disk was subsampled for DNA extraction and stored at -80°C.

### Sodium, biomass concentration, and Spirulina experiments

The dark and anoxic incubation of the cyanobacterial consortium was repeated using dewatered biomass with different concentrations of sodium. Initially, biomass obtained after growth was first dewatered and then gently washed with deionized water to remove the salts. This step was repeated five times to ensure that all the salts were removed. The washed biomass was then separated into three aliquots. Each aliquot was washed with sodium carbonate solution with varied concentrations (0.25M, 0.5M and 1M). Then, approximately 2 grams of wet paste from each aliquot was placed in sterile serum bottles. The headspace in the serum bottles was vacuumed and filled with nitrogen gas up to atmospheric pressure to create anoxic conditions. These serum bottles were then incubated in dark at room temperature for 8 days. Every day two bottles were removed from the incubation and analysed for phycocyanin as described above. Electron microscopy was conducted as described previously [101], without performing ethanol washes and without using gold-sputtered filters.

The dark and anoxic incubation was repeated using a culture of *Arthrospira platensis* (Spirulina). A bulk culture containing 250 g of wet paste (solid concentration 20% w/w) was incubated in dark and anoxic conditions with 1 M sodium carbonate for 12 days. The culture was visually monitored for signs of lysis and phycocyanin release (Fig. S5).

## Acknowledgments

We thank the University of Calgary’s Center for Health Genomics and Informatics for sequencing and informatics services. We thank Carmen Li for help with MiSeq sequencing. We would like to thank Jayne Rattray and Martin Pabst for help with measurements and identification of organic acids.

## Author Contributions

J.Z. extracted DNA and protein, and analyzed metagenomics and metaproteomics data. A.J.P. collected soda lake sediment samples, prepared them for 16S rRNA gene amplicon sequencing, and analyzed data. A.V., T.G., C.D., and A.J.P. performed the incubation experiments. A.K. maintained and ran the mass spectrometer required for metaproteomics. V.K. helped to analyze metagenomes. J.Z., A.V., and M.S. drafted the manuscript with input from all authors. M.S. and H.D. secured funding for this research.

## Data and Materials Availability

All data needed to evaluate the conclusions in the paper are present in the paper and/or the Supplementary Materials. Metaproteomes were deposited to the PRIDE database under accession PXD023504.

Metagenomes can be found in BioSamples SAMN17264972-SAMN17264985 (BioProject: PRJNA377096). MAGs from the study were deposited into the NCBI Whole Genome Shotgun submission database under BioSamples SAMN17266165-SAMN17266224.

