## Supplementary Materials for "A programmed response precedes cell lysis and death in a mat-forming cyanobacterium"

### A naturally occurring lytic bioprocess improves sustainability of phycocyanin production from cyanobacteria

**This file includes:**

Figs. S1 to S6

**Other Supplementary Materials for this manuscript include the following:**

Tables S1 to S2
Supplementary Table 1 - MAG information
Supplementary Table 2 - Metaproteome protein abundances
Supplementary Table 3 – Proteins identified as significantly associated with an incubation fraction

Data S1
Supplementary Data 1 - Predicted proteins and corresponding accessions used in this manuscript from the Ca. P. alkaliphilum MAG

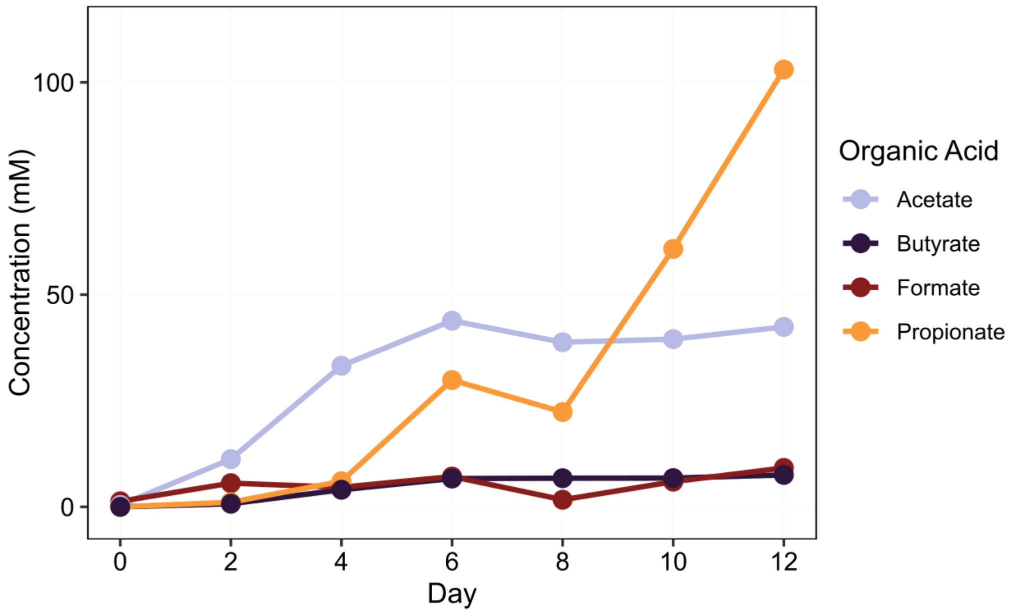


**Fig. S1.**
Concentrations of organic acids in the supernatant fraction during the incubation.

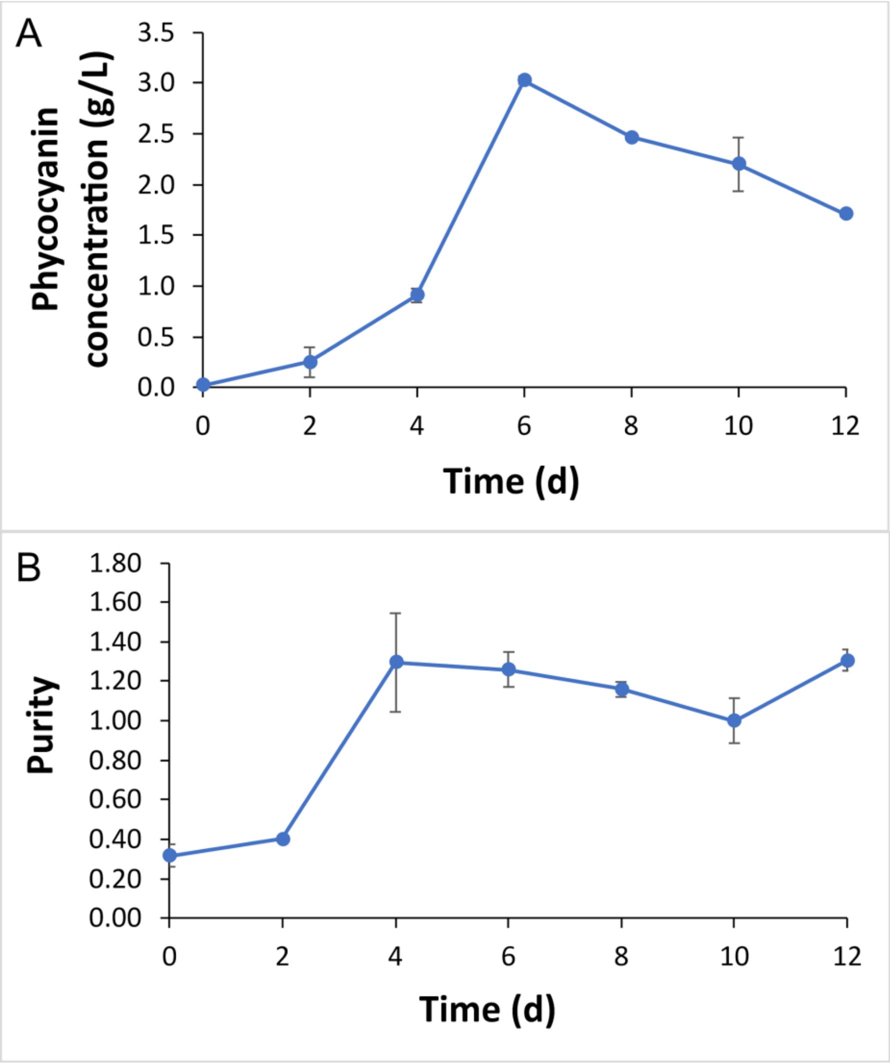


**Fig. S2.**
Concentration and purity of released phycocyanin during the incubation. (A) Phycocyanin concentration measured as absorbance at 620 nm compared to a standard curve of laboratory grade phycocyanin. (B) Purity was measured as the absorbance ratio of 620/280 nm.

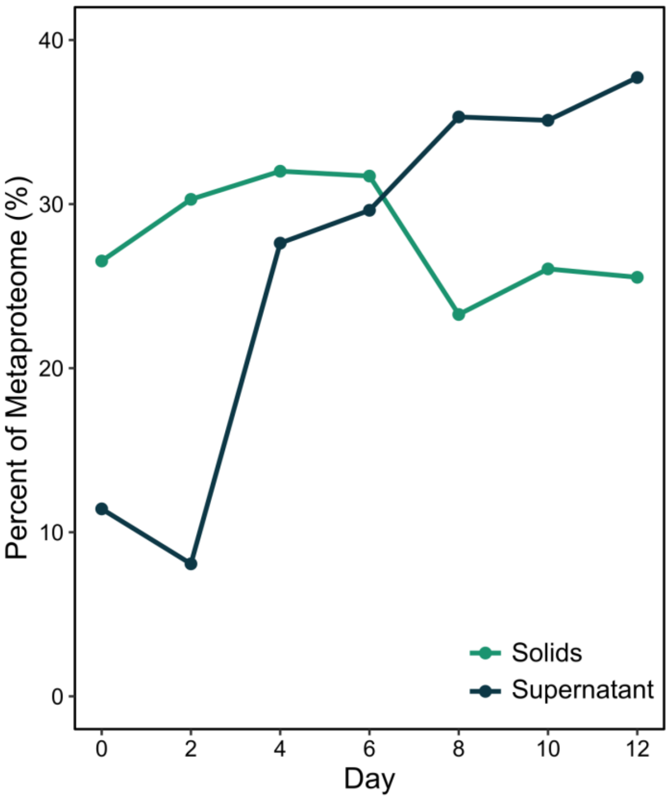


**Fig. S3.**
Relative abundance of *Ca.* P. alkaliphilum pigment proteins in the metaproteome during dark and anoxic incubation. Proteins from the solid fraction are shown in green and proteins from the supernatant fraction are shown in dark blue.

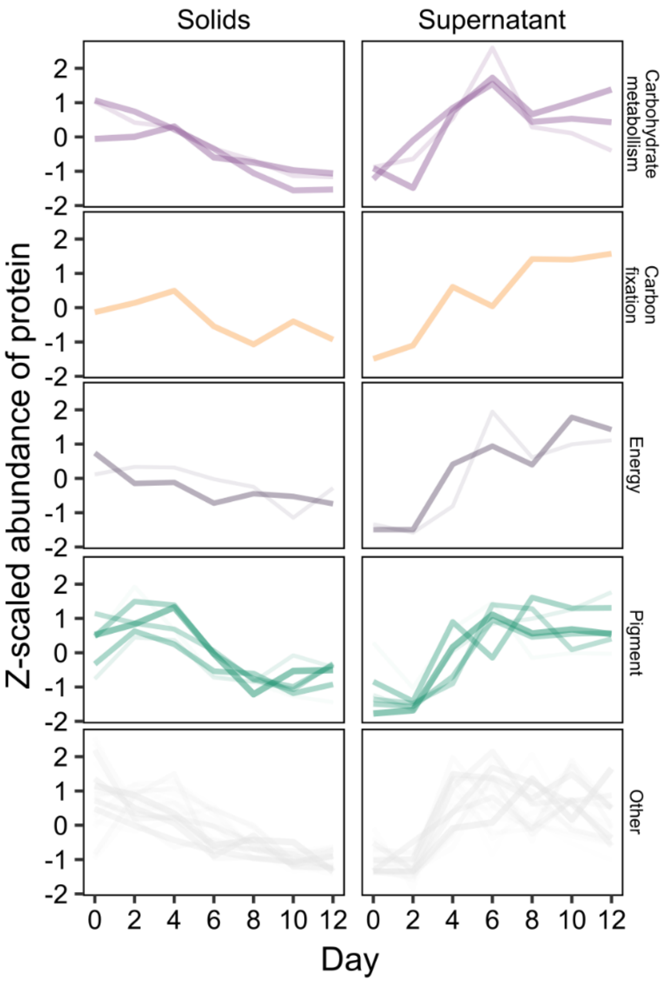


**Fig S4.**
Proteins from functional categories that were significantly associated with both pre-lysis solids and post-lysis supernatant fractions. These proteins were likely intracellular proteins released into the media upon cell lysis. Protein abundances were scaled across samples for visualization purposes. For a complete list of these proteins see Supplementary Table 3.

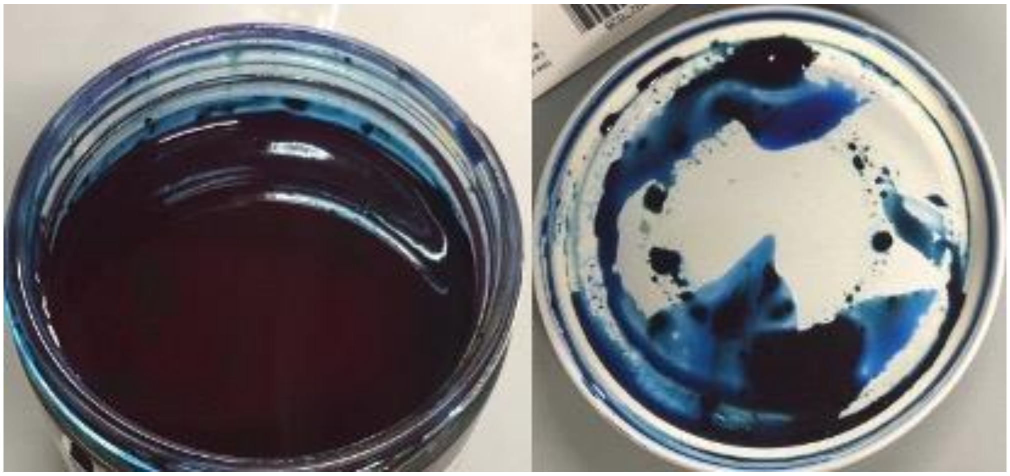

**Fig. S5.**
Image of lysed *Arthrospira platensis* culture after dark and anoxic bulk incubation.

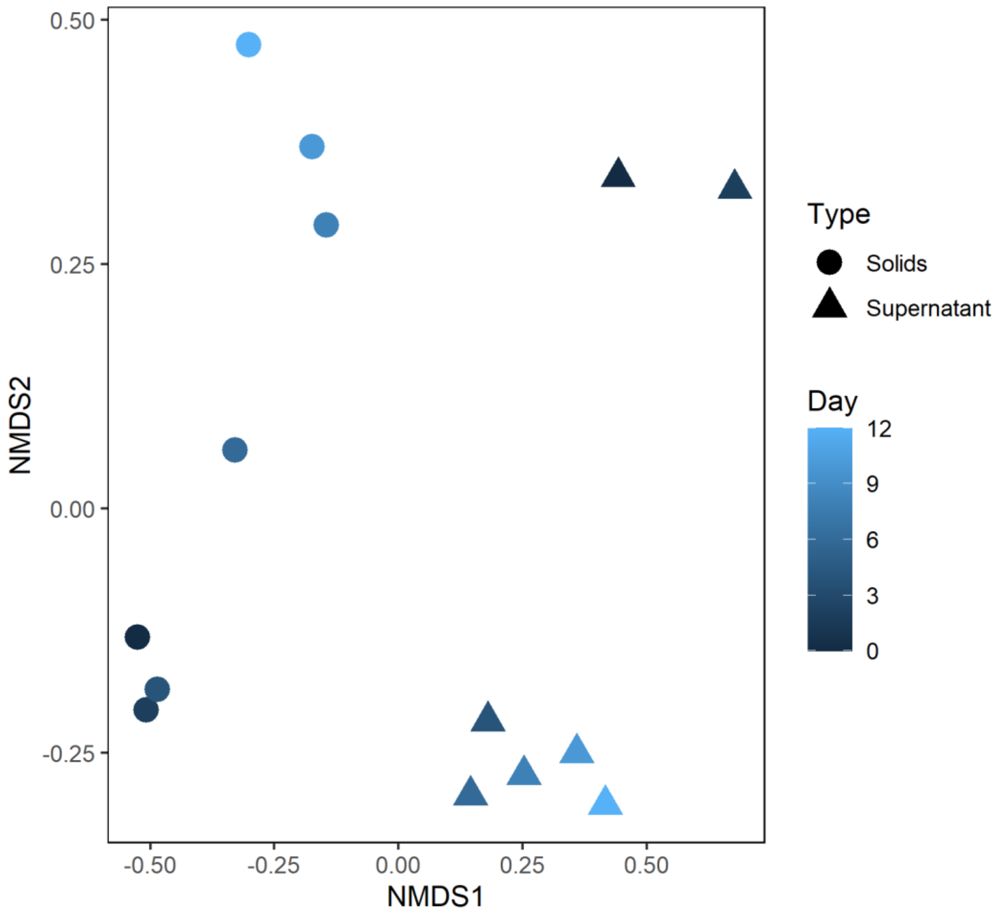


**Fig S6.**
NMDS of incubation samples based on Bray-Curtis dissimilarity of protein abundance. Only proteins with more than 50 PSMs included. Stress = 0.09.

**Table S1. (separate file)**
MAG information. Contains information on all metagenome assembled genomes present in the consortium.

**Table S2. (separate file)**
Metaproteome protein abundances. Contains abundances of all proteins identified in the metaproteomes.

**Table S3. (separate file)**
Proteins identified as significantly associated with an incubation fraction.

**Data S1. (separate file)**
Predicted proteins and corresponding accessions used in this manuscript from the *Ca.* P. alkaliphilum MAG with annotations from MetaErg.
